# Genetic control of the human brain proteome

**DOI:** 10.1101/816652

**Authors:** Chloe Robins, Aliza P. Wingo, Wen Fan, Duc M. Duong, Jacob Meigs, Ekaterina S. Gerasimov, Eric B. Dammer, David J. Cutler, Philip L. De Jager, David A. Bennett, James J. Lah, Allan I. Levey, Nicholas T. Seyfried, Thomas S. Wingo

## Abstract

Alteration of protein abundance and conformation are widely believed to be the hallmark of neurodegenerative diseases. Yet relatively little is known about the genetic variation that controls protein abundance in the healthy human brain. The genetic control of protein abundance is generally thought to parallel that of RNA expression, but there is little direct evidence to support this view. Here, we performed a large-scale protein quantitative trait locus (pQTL) analysis using single nucleotide variants (SNVs) from whole-genome sequencing and tandem mass spectrometry-based proteomic quantification of 12,691 unique proteins (7,901 after quality control) from the dorsolateral prefrontal cortex (dPFC) in 144 cognitively normal individuals. We identified 28,211 pQTLs that were significantly associated with the abundance of 864 proteins. These pQTLs were compared to dPFC expression quantitative trait loci (eQTL) in cognitive normal individuals (n=169; 81 had protein data) and a meta-analysis of dPFC eQTLs (n=1,433). We found that strong pQTLs are generally only weak eQTLs, and that the majority of strong eQTLs are not detectable pQTLs. These results suggest that the genetic control of mRNA and protein abundance may be substantially distinct and suggests inference concerning protein abundance made from mRNA in human brain should be treated with caution.

## Introduction

Proteins have an important role in neurodegenerative disease. Alzheimer’s disease (AD), for instance, is characterized by the abnormal accumulation of amyloid-beta and tau proteins in the brain. In addition to these hallmark proteins, hundreds of other proteins have also been shown to correlate with neurodegenerative disease phenotypes such as rate of decline of cognition and diagnosis of AD (*1, 2*). This suggests that the dysregulation of numerous proteins may contribute to disease.

While the genetic control of mRNA expression in the brain has been well studied (*3–8*), little is known about how genetics influence protein abundance in the brain. Thousands of genetic variants have been reported to associate with variation in mRNA levels, known as expression quantitative trait loci (eQTLs). Oftentimes, these identified genetic effects on mRNA are used to help prioritize candidate causal genes identified by genome-wide association studies (*9*), and these effects are assumed to translate to protein abundance. However, this relationship has yet to be tested with large-scale profiling of brain proteins. A better understanding of the genetic control of protein abundance will help us define the relationships between genetics, gene expression, and proteins, and lead to a better understanding of the molecular changes that underlie neurodegenerative and other brain diseases.

Here, we perform a large-scale investigation into the genetic control of the human brain proteome using high-throughput mass spectrometry-based protein quantification and whole genome sequencing from post-mortem brain samples of the dorsolateral prefrontal cortex of cognitively normal older adults. To understand the relationship between genetics, gene expression, and protein, we also investigate the genetic control of gene expression (i.e. mRNA) in the human brain from the same cohort and among the same individuals with brain proteomes. Finally, we compare the pQTLs identified with eQTLs from a recent meta-analysis of brain expression of over 1,433 brains (*10*). The results of our analyses are available at http://brainqtl.org and serve as a resource for future investigations.

## Methods

Our methods are provided in the supplementary materials section at the end of this document.

## Results

### Demographics

We analyzed genetic, proteomic, and transcriptomic data from 233 ROS/MAP participants, of which 144 have proteomic data, 169 have transcriptomic data, and 81 have both proteomic and transcriptomic data. Table S2 gives the demographic characteristics of these subjects. Participants had a high degree of education (median of 16 years), were all Caucasian, and were 63% women. The age at death ranged from 67 years to 102 years, with a median age at death of 86.5 years.

### Brain QTL Mapping

To identify pQTLs, we analyzed proteomes from the dorsolateral prefrontal cortex and genotyping from WGS of 144 individuals. The genotypes of a total of 2,599,383 SNVs were tested against the abundance of 7,901 proteins. We found that 10.9% (864 of 7,901) of the proteins were associated with a total of 28,211 pQTLs (FDR < 0.05). Of the proteins with a pQTL, 77.7% (671 of 864 proteins) had multiple pQTLs, with an average of 33 pQTLs detected per protein.

To identify eQTLs and pQTLs from the same set of genes, we tested genotypes of 2,082,000 SNVs against the abundance of protein and mRNA from a restricted set of 5,743 genes found in both studies. We found that 10.7% (617 / 5,743) of genes had a pQTL and 14.7% (843 / 5,743) of genes had an eQTL. Fewer pQTLs were identified than eQTLs with a total of 21,034 and 35,064, respectively. Additionally, genes with a pQTL averaged 34 pQTLs per gene compared to 42 eQTLs per gene for genes with an eQTL. Only 199 genes had both a pQTL and an eQTL, which represents 32.3% (199 / 617) of genes with a pQTL and 23.6% (199 / 843) of genes with an eQTL. A total of 3,364 SNVs were identified as both a pQTL and an eQTL (i.e. eQTL/pQTLs) in 95 of the 199 genes with both a pQTL and an eQTL (see Table S3 for the top 10). Thus, only 16.0% (3,364 / 21,034) of all identified pQTLs are eQTLs and 9.6% (3,364 / 35,064) of all identified eQTLs are pQTLs (Figure 1A inset). These results were essentially unchanged when we: 1) limited the analysis to samples with complete proteomic and transcriptomic data (see supplementary materials); 2) used different *R*^2^ thresholds for linkage disequilibrium (see supplementary materials); 3) varied the window of SNVs around the gene (50kb, 100kb, or 500kb; see supplementary materials); 4) used the more stringent Bonferroni significance threshold to define pQTLs and eQTLs (see supplementary materials).

**Fig. 1.**
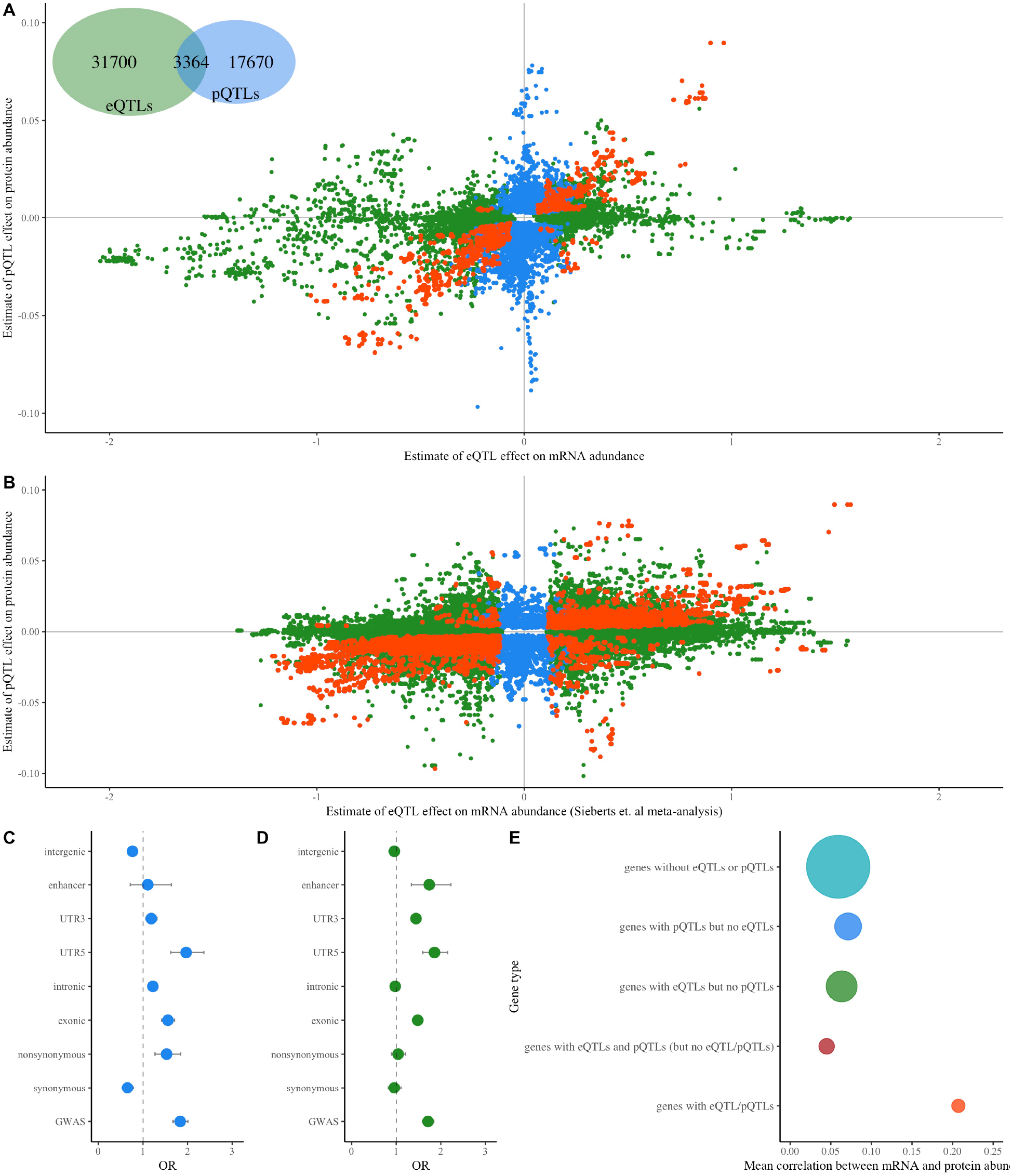
Protein and RNA Quantitative Locus Results. This figure summarizes the direction of effect and genomic annotation for pQTL and eQTL sites. (A) Comparison of eQTL and pQTL estimates. Each point represents one SNV tested against the abundance of the mRNA and protein of a single gene. eQTLs (defined based on False Discovery Rate (FDR) < 0.05) are shown in green, pQTLs (defined based on FDR < 0.05) are shown in blue, and sites that are both an eQTL and a pQTL (i.e. eQTL/pQTLs) are shown in orange. (B) Comparison of Sieberts *et al.* meta-analysis eQTL estimates (N=1,433) and our pQTL estimates. Each point represents one SNV tested against the abundance of the mRNA and protein of a single gene. eQTLs (defined based on FDR < 0.05) are shown in green, pQTLs (defined based on Bonferroni correction < 0.05) are shown in blue, and sites that are both an eQTL and a pQTL (i.e. eQTL/pQTLs) are shown in orange. (C) Results of Fischer’s exact tests for the enrichment of pQTLs. Odds ratio (OR) estimates are shown with 95% confidence intervals. (D) Results of Fischer’s exact tests for the enrichment of eQTLs. OR estimates are shown with 95% confidence intervals. (E) Mean correlation between mRNA and protein abundance for genes without a pQTL or an eQTL, genes with eQTL/pQTLs (i.e. sites that are both an eQTL and a pQTL), genes with pQTLs and eQTLs but no eQTL/pQTLs, genes with pQTLs and no eQTLs, and genes with eQTLs and no pQTLs. The size of the point reflects the relative number of proteins within each gene type.

We tested the reproducibility of our analyses by comparing our eQTLs with those reported previously by (*10*) in a larger sample of 1,433 cognitively normal and cognitively impaired individuals, and saw a high replication rate of 92.8% (see supplementary material). Furthermore, we assessed the relationship between the minor allele frequencies (MAF) of the tested variants and our declaration of pQTLs and eQTLs (see supplementary material). We found that the percentage of the total pQTLs, eQTLs, and eQTL/pQTLs identified slightly increases with variant MAF. The smallest percentage of pQTLs, eQTLs and eQTL/pQTLs identified have a variant MAF between 0.05 and 0.1 (5%-8%), while the largest percentage of pQTLs, eQTLs, and eQTL/pQTLs have a variant MAF between 0.4 to 0.5 (26%-36%). This suggests a slight, but not substantial, dependence of pQTLs and eQTLs on MAF, likely because our power to detect QTLs is higher for variants with higher MAF. Finally, we compared our identified brain pQTLs to previously published pQTLs of human blood proteins (*41–43*) and found very few pQTLs from human brain to also be pQTLs of human blood proteins (3%, see supplementary materials). This suggests that the genetic control of blood and brain proteins are largely distinct.

For each SNV that was identified as either an eQTL or a pQTL, we compared the effect of each variant on mRNA and protein abundance (Figure 1A). To facilitate the comparison, we define strong pQTLs as SNVs with a significant association between genotype and protein abundance (FDR < 0.05) and an effect estimate greater than five standard deviations from the mean effect estimate across all SNV-protein pairs. Similarly, we define strong eQTLs as SNVs with a significant association between genotype and mRNA abundance (FDR < 0.05) and an effect estimate greater than five standard deviations from the mean effect estimate across all SNV-mRNA pairs. Based on these definitions, 12.4% (4,335/ 35,064) of the identified eQTLs are strong eQTLs, 12.0% (2,516 / 21,034) of the identified pQTLs are strong pQTLs, and 18.4% (620 / 3,364) of the sites that are both an eQTL and a pQTL (eQTL/pQTLs) are both a strong eQTL and a strong pQTL. We found that 71.1% (3,083/ 4,335) of strong eQTLs have only weak effects on protein abundance, and that 75.4% (1,896 / 2,516) of strong pQTLs have only weak effects on mRNA abundance. Finally, at sites that are both an eQTL and a pQTL, only 6.0% (203 / 3,364) are strong eQTLs but not strong pQTLs, and 6.5% (218 / 3,364) are strong pQTLs but not strong eQTLs. This means that eQTL/pQTL sites tend to have similar effects on mRNA and protein abundance, with 87.5% (2,943/3,364) having matching effects (strong or weak) on mRNA and protein abundance. Furthermore, 94% (3,200 / 3,364) of these eQTL/pQTLs have effects on mRNA and protein abundance that are in the same direction (Figure 1A).

To assess generalizability, we compared our pQTL results to eQTL results from a large meta-analysis using data from the dPFC of 1,433 samples from four cohorts (*10*). For each SNV that we identified as a pQTL or was identified as an eQTL in the meta-analysis, we compared the effect of each variant on mRNA and protein abundance (Figure 1B). Even with the increase in sample size and power to detect eQTLs, we still see similar patterns between the effects of pQTLs and eQTLs. That is, the majority of strong eQTLs are not strong pQTLs, and vice versa. This suggests that most of the large genetic effects on protein abundance have only a small effect on mRNA levels, and that most of the large genetic effects on mRNA levels have only a small effect on protein abundance.

### Functional analysis of pQTLs and eQTLs

The pQTLs identified were more likely to be located in a genic region (5’ UTR OR: 1.97, FDR adjusted p-value: 1.3×10^−10^, exons OR: 1.56, FDR adjusted p-value: 2.3×10^−18^, introns OR: 1.22, FDR adjusted p-value: 3.0×10^−44^, Figure 1C) and less likely to be located in an intergenic region by Fisher’s exact test (OR: 0.77, FDR adjusted p-value: 8.3×10^−73^). Furthermore, the identified pQTLs are more likely to be non-synonymous nucleotide substitutions (OR: 1.53, FDR adjusted p-value: 6.8×10^−6^), and less likely to be synonymous nucleotide substitutions (OR: 0.65, FDR adjusted p-value: 6.8×10^−6^). The identified eQTLs were more likely to be located in exons (OR: 1.48, FDR adjusted p-value: 2.9×10^−22^), 5’ UTRs (OR: 1.86 FDR adjusted p-value: 1.6×10^−13^), 3’ UTRs (OR: 1.44, FDR adjusted p-value: 3.3×10^−20^), and enhancers (OR: 1.74, FDR adjusted p-value: 7.5 × 10^−5^), and less likely to be located in introns (OR: 0.98, FDR adjusted p-value: 0.04) and in an intergenic region (OR: 0.96, FDR adjusted p-value: 0.00017, Figure 1D). Additionally, identified eQTLs were not found to be significantly more likely to be either non-synonymous (OR: 1.04, FDR adjusted p-value: 0.60) or synonymous nucleotide substitutions (OR: 0.96, FDR adjusted p-value: 0.60). Finally, both pQTLs and eQTLs were found to be significantly enriched in GWAS results (pQTL OR: 1.84, FDR adjusted p-value: 2.12 × 10^−33^; eQTL OR: 1,71, FDR adjusted p-value: 1.42 × 10^−40^).

### Correlation between mRNA and protein abundance

To understand the relationship between mRNA and protein abundance, we examined the correlation between mRNA and protein levels for five sets of genes: 1) genes with sites that are both an eQTL and a pQTL (i.e. eQTL/pQTLs) (n=95); 2) genes with both pQTLs and eQTLs but no eQTL/pQTLs (n=104); 3) genes with pQTLs but no eQTLs (n = 418); 4) genes with eQTLs but no pQTLs (n = 644); 5) genes without any pQTLs or eQTLs (n = 4,482). Remarkably, genes with sites that are both eQTLs and pQTLs (eQTL/pQTLs) have the highest average correlation between mRNA and protein level (0.21), while all other genes have notably lower average correlations (0.04 to 0.07) (Figure 1E). Additionally, we found the distributions of correlations between mRNA and protein level to be significantly different between the genes with eQTL/pQTLs, genes with both pQTLs and eQTLs but no eQTL/pQTLs, genes with pQTLs but no eQTLs, genes with eQTLs but no pQTLs, and genes with no eQTLs or pQTLs (Kruskal-Wallis test, chi-square = 82.6, p < 2.2×10^−16^). Specifically, the median correlation between mRNA and protein levels was found to be significantly higher for genes with eQTL/pQTLs than genes with both pQTLs and eQTLs but not eQTL/pQTLs (0.21 vs. 0.04, Wilcoxon test, p-value:6.4×10^−11^), genes with only pQTLs (0.21 vs. 0.07, Wilcoxon test, p-value: 1.3×10^−10^), genes with only eQTLs (0.21 vs. 0.06, Wilcoxon test, p-value: 1.4×10^−13^), and genes without eQTLs or pQTLs (0.21 vs. 0.06, Wilcoxon test, p-value: 3.3×10^−16^).

### Brain QTL Resource

Our pQTL and eQTL results are available online at http://brainqtl.org. This website visually displays the results of our pQTL and eQTL analyses and provides summary statistics on individual variants tested.

## Discussion

We performed the first unbiased large-scale investigation into the genetic control of the human brain proteome and presented evidence for thousands of pQTLs influencing abundance of hundreds of brain proteins. Our most striking result was that the majority of strong pQTLs are weak eQTLs, and vice versa. Additionally, only a small minority of protein-coding genes have an SNV that is appreciably associated with mRNA and protein abundances and can be defined as both a pQTL and an eQTL. This result is consistent with previous work in other tissues that also found few sites that are both a pQTL and an eQTL (*41, 44, 45*).

We found the small number of genes that have a SNV that is both a pQTL and an eQTL (i.e. an eQTL/pQTL) to have higher correlations between mRNA and protein abundance than genes with only pQTLs or eQTLs alone. This suggests that protein abundance may be limited by transcript availability for genes with eQTL/pQTLs, and controlled by post-transcriptional processes such as miRNA, localization (e.g. membrane, intra- or extracellular), translational regulation, or post-translational modifications for genes with pQTLs but no eQTLs or only weak eQTLs. The depletion and enrichment of pQTLs for non-synonymous and synonymous sites, respectively, also suggests translational regulation of protein abundance. Together these results suggest that caution is needed when inferring the effect of an eQTL on protein abundance in the human brain.

Both pQTLs and eQTLs were found to be enriched for SNVs with a reported GWAS association. Previous studies have reported an enrichment of eQTLs in SNVs associated with complex traits (*46*) as well as an enrichment of disease susceptibility SNVs in brain eQTLs (*3*). Our pQTL and eQTL results suggest that pQTLs and eQTLs are more likely to have a role in disease susceptibility, but the mechanisms by which pQTLs and eQTLs contribute to disease susceptibility likely differs.

We identified 1.7 times more eQTLs than pQTLs. This is consistent with work in other tissues from both humans and mice that also found fewer pQTLs than eQTLs (*41, 42, 44, 45, 47*). The difference in the number of pQTLs and eQTLs may be due to greater tolerance for differences in mRNA abundance (*48*). Furthermore, the protein lifecycle has considerable variability and includes numerous contributing processes beyond transcription. In many cases, the effect of genetic variants on protein abundance may be lessened by post-transcriptional mechanisms (*44*). This is compatible with both our observation of more genetic variants associated with mRNA abundance than protein abundance and our observation of differing effect sizes for the genetic control of protein and mRNA abundance.

Our results should be interpreted with respect to the strengths and limitations of this study. To our knowledge, the brain proteomic sequencing performed here is among the deepest thus far profiled with a total of 12,691 unique proteins (12,415 detected genes). This reflects about 79% of all expressed transcripts in the human brain. To achieve throughput and proteomic depth of sequencing, we used TMT isobaric labeling coupled with high-pH offline fractionation following well-established protocols (*26*). A limitation of the proteomic data was the use of MS2 acquisition, which can suffer from the presence of co-isolated and co-fragmented interfering ions that can obscure quantification (*49*). However, high-pH offline fractionation largely mitigates this issue (*26*). Another potential limitation is that the observed differences in the number of pQTLs and eQTLs may be in part due to technical differences in the proteomic and transcriptomic profiling methods. However, we note that a previous plasma-based pQTL analysis made a similar observation in blood and used the SOMAscan technology, which differs from the mass spectrometry-based proteomics used here (*41*). Another potential limitation is that the number of pQTLs and eQTLs may be limited by the power to detect them, as suggested by our sensitivity analysis (see supplementary materials). However, our study was still sufficiently powered to detect thousands of pQTLs and eQTLs, and these potential power issues do not appear to influence our main conclusion regarding the control of brain protein and RNA expression (see supplementary materials). Another potential limitation of this study is that both the gene expression and protein data were generated from bulk tissue that is composed of a mixture of cell types. We estimated cell type proportions for both data types, but found the inclusion of estimated cell type proportions in our analyses to increase test-statistic inflation (see supplementary materials). This suggests that available methods for cell type deconvolution are not sufficiently able to remove confounding by cell type for human brain proteomic data. Despite these limitations, our study is the first to examine genetic control of human brain proteins and reveals key differences in pQTLs versus eQTLs in the human brain and provides a web-based resource to enable researchers to explore the genetic control of the human brain proteome.

## Acknowledgments

We thank the participants of the ROS/MAP studies and Thanneer Perumal, PhD for creating the estimated cell type proportions from RNA-sequencing data.

## Author contributions

Conceptualization (A.P.W, C.R., D.J.C., T.S.W.), Data curation (C.R., D.M.D., E.B.D., N.T.S.), Formal Analysis (A.P.W, C.R., D.J.C., T.S.W., W.F.), Funding acquisition (A.I.L, A.P.W, D.A.B., N.T.S., P.L.D., T.S.W.), Investigation (D.M.D., E.B.D., N.T.S.), Project administration (A.I.L, A.P.W, D.A.B., J.J.L., N.T.S., T.S.W.), Resources (D.A.B., P.L.D.), Software (J.M., T.S.W.), Supervision (A.P.W, T.S.W.), Validation (C.R., D.M.D., E.B.D., E.S.G., W.F.), Visualization (C.R., J.M.), Writing – original draft (C.R.), Writing – review & editing (All authors).

## Data and materials availability

Study data were provided by the Rush Alzheimer’s Disease Center, Rush University Medical Center, Chicago. The results published here are in whole or in part based on data obtained from the AMP-AD Knowledge Portal (doi:10.7303/syn2580853). RADC Resource Sharing Hub: http://www.radc.rush.edu/. Synapse website: https://www.synapse.org. Our pQTL and eQTL results are available for download and visualization at: http://brainqtl.org.

## Supplementary Materials

### Materials and Methods

#### Study subjects

The subjects in this study are participants of the Religious Orders Study (ROS) and the Memory and Aging Project (MAP). ROS and MAP are longitudinal cohort studies of Alzheimer’s disease and aging maintained by investigators at the Rush Alzheimer’s Disease Center in Chicago, IL (*11–13*). Both studies recruit participants without known dementia at baseline and follow them annually using detailed clinical evaluation. ROS recruits individuals from catholic religious orders from across the USA, while MAP recruits individuals from retirement communities as well as individual home visits in the Chicago, IL area. Participants in each study undergo annual medical, neurological, and neuropsychiatric assessments from enrollment to death, and neuropathologic evaluations at autopsy. Participants provided informed consent, signed an Anatomic Gift Act, and repository consent to allow their data and biospecimens to be repurposed. The studies were approved by an Institutional Review Board of Rush University Medical Center.

#### Clinical Diagnoses

For each ROS/MAP participant, a clinical diagnosis of dementia is rendered annually and at the time of death. The diagnosis rendered at death is based on all available clinical data and is given by a neurologist who is blinded to all postmortem data using the National Institute of Neurological and Communicative Disorders and Stroke and the Alzheimer’s Disease and Related Disorders Association guidelines (*14*). Case conferences including two neurologists and a neuropsychologist were used for consensus, as necessary, for select cases. Diagnoses of dementia status was coded as no cognitive impairment (NCI), mild cognitive impairment (MCI), or Alzheimer’s dementia (AD). Here, we restricted the analyses to those with NCI at death to investigate the genetic control of the normal human brain proteome and transcriptome. ROS/MAP resources can be requested at www.radc.rush.edu.

#### Genetic data

Genotype data was generated from whole genome sequencing (WGS) of DNA that was extracted from cryopreserved peripheral blood mononuclear cells or frozen dorsolateral prefrontal cortex (dPFC) of ROS/MAP subjects. WGS was performed as described in detail by De Jager et al. (*15*) and is available via Synapse (ID: syn10901595). Briefly, libraries were constructed using the KAPA Hyper Library Preparation Kit per the manufacturer’s protocol and sequenced on an Illumina HiSeq X sequencer (v2.5 chemistry) using 150bp paired-end reads. Reads were aligned to the GRCh37 human reference genome using Burrows-Wheeler Aligner (BWA-MEM v0.7.8) (*16*) and processed using the GATK best-practices workflow, which includes marking duplicate reads by Picard tools v1.83, local realignment around indels, and base quality score recalibration by Genome Analysis Toolkit (GATK v3.4.0) (*17, 18*). A multi-sample genomic variant call format (gVCF) was generated by merging results of HaplotypeCaller on each sample individually in gVCF mode (GATKv3.4.0) and batches of gVCF were merged into gVCFs processed by a joint genotyping step (GATK v3.2.2).

Annotation of the multi-sample VCF (n=1,196) was performed using Bystro (*19*) and supplemented by the Broad’s ChromHMM annotation of dPFC tissue (*20, 21*). A total of 1,133 samples passed all quality control measures and 63 samples were excluded for one or more of the following reasons. Samples with greater than five standard deviations for *θ*, silent:replacement sites, and transition:transversion ratio were excluded (n=7), and samples with greater than three standard deviations for genotype missingness, heterozygosity, or homozygosity were excluded (n=14). Samples that were discordant for sex based on heterozygosity of the X chromosome were also excluded (n=7). Cryptically related or duplicate samples were identified by identity-by-state sharing using PLINK (*22*) and removed (n=31). Unlinked ancestrally informative markers were used to infer eigenvectors for principal-component analysis using EIGENSTRAT (*23*) and over six standard deviation outliers (n=1) were removed. Before analysis, we also removed all non-SNVs (i.e. insertions and deletions), SNVs outside of Hady-Weinberg equilibrium, SNVs with missing data for over 10% of samples, and SNVs with a minor allele frequency less than 0.05.

#### Protein Abundance by Tandem MS-based Proteomics

Protein abundance from cortical microdissections of dPFC (Broadman area 9) of ROS/MAP subjects was generated using tandem mass tag (TMT) isobaric labeling mass spectrometry methods for protein identification and quantification. Tissue homogenization was performed as described by (*24*), followed by protein digestion. For protein digestion, 100 μg of each sample was reduced with 1 mM dithiothreitol (DTT) at room temperature (RT) for 30 min, followed by 5 mM iodoacetamide (IAA) alkylation in the dark for another 30 min and overnight digestion with Lysyl endopeptidase (Wako) at 1:100 (w/w). Subsequently, samples were diluted 7-fold with 50 mM ammonium bicarbonate (AmBic) and digested with 1:50 (w/w) Trypsin (Promega) for another 16 h. The peptide solutions were acidified to a final concentration of 1% (vol/vol) formic acid (FA) and 0.1% (vol/vol) triflouroacetic acid (TFA), desalted with a 30 mg HLB column (Oasis). An equal amount of protein from each sample was aliquoted and digested in parallel to serve as the global pooled internal standard (GIS) in each TMT batch.

Prior to TMT labeling, all samples were randomized into 50 batches (8 samples per batch) based on age at death, sex, post-mortem interval, diagnosis, and measured neuropathologies. Peptides from each individual sample (*n*=400) and the GIS (*n*=100) were labeled using the TMT 10-plex kit (ThermoFisher). In each batch, TMT channels 126 and 131 were used to label GIS standards, while the 8 middle TMT channels were reserved for individual samples following randomization. The TMT labeling was performed as described by (*24, 25*), followed by high-pH fractionation performed as described by (*26*) with slight modification. Dried samples were re-suspended in high pH loading buffer (0.07% vol/vol NH_4_OH; 0.045% vol/vol formic acid, 2% vol/vol acetonitrile) and loaded onto an Agilent ZORBAX 300Extend-C18 column (2.1mm x 150 mm with 3.5 µm beads). An Agilent 1100 HPLC system was used to carry out the fractionation. A total of 96 individual fractions were collected across the gradient and pooled into 24 fractions and dried.

All fractions were resuspended in equal volume of loading buffer (0.1% formic acid, 0.03% trifluoroacetic acid, 1% acetonitrile) and analyzed by liquid chromatography coupled to mass spectrometry as described by (*27*) with slight modifications. Peptide eluents were separated on a self-packed C18 (1.9 um Dr. Maisch, Germany) fused silica column (25 cm × 75 μM internal diameter (ID); New Objective, Woburn, MA) by a Dionex UltiMate 3000 RSLCnano liquid chromatography system (ThermoFisher Scientific) and monitored on an Orbitrap Fusion mass spectrometer (ThermoFisher Scientific). Sample elution was performed over a 180-min gradient with flow rate at 225 nL/min. The gradient goes from 3% to 7% buffer B in 5 mins, from 7% to 30% over 140 mins, from 30% to 60% in 5 mins, 60% to 99% in 2 mins, kept at 99% for 8 min and back to 1% for additional 20 min to equilibrate the column. The mass spectrometer was set to acquire in data dependent mode using the top speed workflow with a cycle time of three seconds. Each cycle consisted of one full scan followed by as many MS/MS (MS2) scans that could fit within the time window. The full scan (MS1) was performed with an m/z range of 350-1500 at 120,000 resolution (at 200 m/z) with AGC (automatic gain control) set at 4×10^5^ and maximum injection time of 50 msec. The most intense ions were selected for higher energy collision-induced dissociation (HCD) at 38% collision energy with an isolation of 0.7 m/z, a resolution of 30,000 and AGC setting of 5×10^4^ and a maximum injection time of 100 msec. Five of the 50 TMT batches were run on the Orbitrap Fusion mass spectrometer using the SPS-MS3 method as previously described by (*24*).

All raw files were analyzed using the Proteome Discoverer suite (version 2.3 ThermoFisher Scientific). MS2 spectra were searched against the canonical UniProtKB Human proteome database (downloaded February 2019 with 20,338 total sequences). The Sequest HT search engine was used and parameters were specified with the following: fully tryptic specificity, maximum of two missed cleavages, minimum peptide length of six, fixed modifications for TMT tags on lysine residues and peptide N-termini (+229.162932 Da) and carbamidomethylation of cysteine residues (+57.02146 Da), variable modifications for oxidation of methionine residues (+15.99492 Da) and deamidation of asparagine and glutamine (+0.984 Da), precursor mass tolerance of 20 ppm, and a fragment mass tolerance of 0.05 Da for MS2 spectra collected in the Orbitrap (0.5 Da for the MS2 from the SPS-MS3 batches). Percolator was used to filter peptide spectral matches (PSM) and peptides to a false discovery rate (FDR) of less than 1%. Following spectral assignment, peptides were assembled into proteins and were further filtered based on the combined probabilities of their constituent peptides to a final FDR of 1%. In cases of redundancy, shared peptides were assigned to the protein sequence in adherence with the principles of parsimony. Reporter ions were quantified from MS2 or MS3 scans using an integration tolerance of 20 ppm with the most confident centroid setting.

The GIS channels 126 and 131 in each batch served as technical replicates. We compared the measured abundance of each protein from the two GIS, and found the measured proteome between the two GIS to be over 99% correlated for all batches (Figure S1). As a quality control measure, we removed protein abundance measurements with low correlations between the two GIS (outside the 95% confidence interval) in each batch before further analysis.

For this study, we included only cognitively normal subjects based on the clinical diagnosis of cognitive status rendered at death. To ensure the analysis of high-quality data we: 1) excluded proteins with missing data for over 50% of samples; 2) scaled each abundance value by a sample-specific total protein abundance measure to remove the effects of loading differences; 3) transformed the data to the log_2_ scale. Outlier samples were then identified and removed through iterative principal component analysis. In each iteration we removed samples more than four standard deviations from the mean of the first or second principal component and then re-calculated all the principal components. Following outlier removal, we again removed proteins with missing data for over 50% of the samples. Finally, the abundance of each protein was residualized using a linear regression model to remove the effects of sex, age at death, post-mortem interval, study, batch, and MS2 versus MS3 reporter quantitation mode.

#### Gene Expression by RNA Sequencing

Gene expression was measured from the dPFC (Broadman area 46) by De Jager et al. (*15*). Briefly, RNA was extracted from cortically dissected sections of dPFC grey matter and samples with RNA integrity numbers (RIN) over 5 were used to prepare RNA-Seq libraries using strand-specific dUTP method with poly-A selection (*28, 29*) using the Illumina HiSeq with 101-bp paired-end reads to a target coverage of 50 million reads per library. Raw RNA-Seq reads were aligned to a GRCh38 reference genome and gene counts were computed using STAR (*30*) as described in reference (*31*). We obtained RNA-Seq data from synapse (ID: syn17010685) and performed the following quality control measures in a subset of individuals with normal cognition defined by a clinical diagnosis of no cognitive impairment rendered at death. We excluded these items: 1) non-Caucasian samples; 2) outlier samples via the GTex expression outlier test (D-statistic below 0.9) (*3, 32*); 3) genes with < 1 cpm in over 50% of samples. Subsequently, the filtered data was normalized using the varianceStabilizingTransformation function from the DESeq2 R package, which log_2_ transforms counts, normalizes for library size, and transforms counts to be approximately homoscedastic (*33*). Lastly, the residual expression for each gene was estimated using linear regression to remove the effects of sex, sequencing batch, age at death, post-mortem interval, RIN, and study.

#### Estimation of confounders

To reduce confounding due to population structure, the first ten principal components derived from principal component analysis of the WGS data were added as model covariates in all relevant analyses. All ten of these principal components had significant Tracey-Widom statistics (p-value < 0.05).

#### Statistical Analyses

To identify genetic variants associated with protein abundance in the brain, we used linear regression to model protein abundance as a function of genotype. We reduced our computational and testing burden by investigating only the proximal genetic effects of common SNV variants by testing only SNVs within 100 Kb of each protein coding gene with a minor allele frequency (MAF) over 5%. The location of each protein coding gene was defined by the knownGene table (GRCh37/hg19 assembly) from the University of California, Santa Cruz (UCSC) table browser (*34*). For each SNV-protein pair, we regressed genotype against protein abundance, assuming additive genetic effects and including the first ten genetic principal components as covariates. We also performed analyses that included cell type proportions and additional unmeasured confounders estimated using the PEER software package in R (*35, 36*) as covariates, but we found their inclusion as covariates to increase *λ* (*37*), an estimate of test statistic inflation (see supplementary materials and table S1). For these analyses, the proportion of neurons, astrocytes, microglia, and oligodendrocytes were estimated for each sample using a modified CIBERSORT (*38*) pipeline with proteomic profiles from isolated mouse brain cell types as the reference (*39*). SNVs where genotype was significantly associated with protein abundance after False Discovery Rate (FDR) correction for multiple comparisons were declared protein quantitative trait loci (pQTLs; FDR < 0.05).

For each protein-coding gene, we also identified genetic variants associated with gene expression in the brain (i.e. mRNA expression). We used the same methods that were used to identify genetic variants associated with protein abundance. That is, we used linear regression to model mRNA abundance as a function of genotype for common SNVs (MAF > 0.05) within 100 kb of each protein-coding gene. To be able to compare SNVs associated with mRNA expression to those associated with protein abundance, we restricted our expression analyses to mRNA transcripts of genes that code for proteins present in our pQTL analyses. The location of each gene was defined by the Ensembl stable gene (EnsGene) table (GRCh37/hg19 assembly) from the University of California, Santa Cruz (UCSC) table browser. This table allowed us to match each EnsGene in our GRCh38-aligned RNAseq dataset with its GRCh37 location. For each SNV-mRNA pair, we regressed genotype against mRNA abundance, assuming additive genetic effects and including the first ten genetic principal components. We also performed analyses that included cell type proportions and additional unmeasured confounders estimated using the PEER software package in R (*35, 36*) as covariates, but found their inclusion as covariates to increase *λ* (*37*) (see supplementary materials and table S1). For these analyses, the proportion of neurons, astrocytes, microglia, and oligodendrocytes were estimated for each sample using the CIBERSORT (*38*) pipeline with single-cell RNA-sequencing expression profiles from isolated human brain cell types (*40*). SNVs where genotype is significantly associated with gene expression after FDR correction for multiple comparisons were declared expression quantitative trait loci (eQTLs; FDR < 0.05).

We assessed whether the identified pQTLs and eQTLs are more likely than other sites to be a published GWAS result, a synonymous or nonsynonymous nucleotide substitution, or in a particular genic location (i.e. exons, introns, 5’ UTRs, 3’UTRs, intergenic regions, enhancers) by testing the overlap between the sets of pQTL and eQTLs sites and the sets of sites reported in NHGRI-EBI Catalog of Published genome-wide association studies (Downloaded July 2019), and we annotated to each substitution type and genic location using Fisher’s exact tests.

### Supplementary Text

#### Estimation and analysis of confounders

To avoid potential confounding due to differences in the cellular composition of the tissue samples, we estimated the cellular composition of each sample for both the transcriptomic and proteomic data and included these estimates as covariates in all relevant analyses. For the transcriptomic data, the proportion of neurons, astrocytes, microglia, and oligodendrocytes were estimated for each sample using the CIBERSORT (*38*) pipeline with single-cell RNA-sequencing expression profiles from isolated human brain cell types (*40*). For each individual proteome, the proportion of neurons, astrocytes, microglia, and oligodendrocytes were estimated using a modified CIBERSORT (*38*) pipeline with proteomic profiles from isolated mouse brain cell types as the reference (*39*). For the proteomic data, the CIBERSORT pipeline was modified to allow for a negative relationship between cell type and the protein levels. This change produced estimates of cell type heterogeneity similar to those estimated with the unmodified CIBERSORT pipeline and RNAseq data.

Unknown confounders in the proteomic and transcriptomic data were estimated using the PEER software package in R (*35, 36*). This software uses factor analysis to identify confounders in the data that were not measured in the study. We estimated ten unknown confounders while protecting the first ten genetic principal components and the estimated cell type proportions. We included the first five estimated unknown confounders as covariates in relevant analyses.

To select the covariates to include in our regression models, we assessed the ability of genetic principal components, estimated cell type proportions, and estimated unknown confounders to reduce inflation in our pQTL and eQTL analyses. To do this we ran the pQTL and eQTL analyses on chromosome 1 with different combinations of covariates, and assessed inflation with the inflation factor *λ*, which estimates the amount of inflation by comparing the distribution of the observed test statistics to that expected under the null hypothesis of no effect (Table S1) (*37*). We found *λ* to increase or remain unchanged when estimated cell type proportions and hidden factors were included as covariates in the models. For this reason, we chose to only include the first ten gene principal components as covariates in all our pQTL and eQTL analyses.

#### QTLs among individuals with paired transcriptomic and proteomic data

The sample sizes and missingness structure differ between the pQTL (n=144) and eQTL (n=169) analyses. The missingness structure differs between the proteomic and transcriptomic data due to the batch-specific measurement of proteins in TMT proteomic experiments, which measure proteins for either all or none of the samples in a batch. Therefore, the number of samples with data differs for each protein, and ranges from 69 samples to 144 samples (Figure S2). The transcriptomic data, on the other hand, has data for same number of samples (N=169) for each gene. As a result of these technical and sample size differences between the proteomic and transcriptomic data, there are slight differences in power between our proteomic and transcriptomic analyses. To investigate whether differences in our pQTL and eQTL results are due to differences in sample size or population, we performed pQTL and eQTL analyses using the subset of 81 samples that have both proteomic and transcriptomic data. Furthermore, we subset the tested proteins and corresponding protein-coding genes to those with 81 samples for both proteomic and transcriptomic data. In this analysis, we tested the genotypes of 1,495,066 SNPs against the abundance of 3,579 proteins and 3,583 mRNAs of the corresponding protein-coding genes. We found 6,149 pQTLs for 230 different proteins (FDR < 0.05) and 3,023 eQTLs for 96 different mRNAs of protein-coding genes (FDR < 0.05). Additionally, only 397 SNVs in 14 protein coding genes are both an eQTL and a pQTL (eQTL/pQTLs).

We compared the effect of each variant on mRNA and protein abundance (Figure S3), and found that 52% (3,210 /6,149) of the identified pQTLs are strong pQTLs, 6% (190 / 3,023) of the identified eQTLs are strong eQTLs, and none of the sites that are both an eQTL and a pQTL (eQTL/pQTLs) are both a strong eQTL and strong pQTL. The sets of strong eQTLs and pQTLs are completely distinct. That is, all of the strong eQTLs are weak pQTLs, and all of the strong pQTLs are weak eQTLs.

The small number of eQTL/pQTLs identified in this analysis matches that seen in all our analyses with larger sample sizes. Furthermore, in all analyses we see that strong pQTLs are usually weak eQTLs, and vice versa. This indicates that the differences we observe in the genetic variants controlling the proteome and transcriptome are not entirely due to differences in the numbers of samples with proteomic and transcriptomic data.

The relationship between the number of eQTLs and pQTLs identified in this analysis (i.e. more pQTLs than eQTLs) is opposite that seen in our main analyses. This may indicate that the observed differences between the numbers of pQTLs and eQTLs could be due to differences in sample size and power, and suggest that many more pQTLs exist than have been identified. This result could also be due to larger pQTL effect sizes in the subset of proteins measured in all samples than in the complete proteome. Proteins that are easily measured with TMT mass spectrometry methods may be of a certain class of protein that have pQTLs of larger effect than is seen across the proteome. Larger samples sizes are necessary to be able to disentangle this observation.

#### Linkage Disequilibrium Threshold Influence on QTLs

Many of the pQTLs we detected are in linkage disequilibrium (LD), which increased our testing burden and lowered our p-value threshold for significance; however, pruning sites based on LD may lead us to obscuring functional genomic relationships. To understand how LD affects our ability to detect proteins with pQTLs, we preformed FDR correction for multiple comparisons based on the number of tests performed after removing SNPs in complete LD (r^2^ =1). We tested the genotypes of 1,114,116 SNPs against the abundance of 7,901 proteins. We found 8,323 pQTLs for 755 different proteins (FDR < 0.05). Since the percentage of proteins with a pQTL decreased from 10.9% to 9.6%, we elected to not prune SNVs that are in LD in any of our analyses.

#### Influence of Window Size on QTLs

To understand how our results are influenced by our choice of window size, we investigated eQTLs and pQTLs within 50 kb, 100 kb, and 500 kb of the TSS of each protein-coding gene. The number of sites and tests substantially increased with each increasing window size. We tested 1,567,675 SNPs in the 50kb window, 2,082,000 SNPs in the 100 kb window, and 3,978,062 SNPs in the 500kb window. After correcting for the number of tests performed for each window size, we found 18,503 pQTLs (FDR < 0.05) and 28,065 eQTLs (FDR < 0.05) in the 50kb window, 21,033 pQTLs (FDR < 0.05) and 35,064 eQTLs (FDR < 0.05) in the 100kb window, and 17,887 pQTLs (FDR < 0.05) and 30,804 eQTLs (FDR < 0.05) in the 500 kb window. In the 500kb window results, 61% of the pQTLs and 56% of the eQTLs are located within 50 kb of the TSS of the associated protein-coding gene, and 76% of the pQTLs and 76% of the eQTLs are located within 100 kb of the TSS of the associated protein-coding gene. By using the 100 kb window we can reduce the number of tests performed and still capture the majority of pQTLs and eQTLs.

#### QTL mapping using Bonferroni significance thresholds

To assess the robustness of our results, we preformed all QTL mapping analyses using the more stringent Bonferroni significance threshold to define pQTLs and eQTLs. As expected using a more stringent significance threshold resulted in fewer sites and genes identified; however, it did not change the relationship between the identified pQTLs and eQTLs or overall nature of the sites identified compared to the results from the FDR adjusted analysis presented in the main text.

A total of 2,082,000 SNVs were tested against the abundance of protein and mRNA from 5,739 genes. We found that 1.3% (76 / 5,739) of genes had a pQTL and 1.9% (112 / 5,739) of genes had an eQTL at Bonferroni significance threshold. Fewer pQTLs were identified than eQTLs with a total of 1,981 and 7,355, respectively. Additionally, genes with a pQTL averaged 26 pQTLs per gene compared to 65 eQTLs per gene for genes with an eQTL. Only 15 genes had both a pQTL and an eQTL, which represents 19.7% (15 / 76) of genes with a pQTL and 13.4% (15 / 112) of genes with an eQTL. A total of 332 SNVs were identified as both pQTLs and eQTLs in 12 of the 15 genes with both a pQTL and an eQTL (Table S4). Thus, only 16.8% (332 / 1981) of all identified pQTLs are eQTLs and 4.5% (332 / 7355) of all identified eQTLs are pQTLs (Figure S4A inset). The relationships between the numbers of eQTLs, pQTLs, and eQTL/pQTLs identified in this analysis mirror the relationships seen in our previous analyses using less conservative FDR significance thresholds to define pQTLs and eQTLs. In all analyses we see a greater number of eQTLs than pQTLs, and that the majority of eQTLs are not pQTLs, and vice versa.

For each SNV that was identified as either an eQTL or a pQTL using a Bonferroni significance threshold, we compared the effect of each variant on mRNA and protein abundance (Figure S4 A). This revealed similar results to the analysis using FDR adjusted p-values presented in the main text. Using the definition of a strong QTL as a site associated with changing expression or protein abundance greater than two standard deviations from the mean, we found that most eQTLs are not strong pQTLs and vice versa. For the sites that are associated with both mRNA and protein abundance (i.e. eQTL/pQTLs), the majority of the effects on mRNA and protein abundance are in the same direction (94%, 313/332). To help understand if these results are generalizable, we compared the pQTLs identified at Bonferroni adjusted threshold to the results of eQTL meta-analysis of brain using the dPFC of 1,433 samples from four cohorts (*10*). For each SNV that we identified as a pQTL (defined by Bonferroni adjusted p-value threshold) or was identified as an eQTL in the meta-analysis, we compared the effect of each variant on mRNA and protein abundance (Figure S4 B). Similar to what was found using FDR threshold to define pQTLs, we found the majority of strong eQTLs are not pQTLs, and that strong pQTLs are usually weak eQTLs.

The location of sites defined by either Bonferroni or FDR significance thresholds were, essentially, the same. The Bonferroni-defined pQTLs were more likely to be located in a genic region (5’ UTR OR: 3.2 adjusted p-value: 4.6×10^−5^, exons OR: 2.8, adjusted p-value: 1.2×10^−13^, introns OR: 1.3, adjusted p-value: 2.0×10^−8^, Figure S4 C) and less likely to be located in an intergenic region by Fisher’s exact test (OR: 0.6, FDR adjusted p-value: 2.9×10^−18^, Figure S4 C). Furthermore, they were more likely to be non-synonymous nucleotide substitutions (OR: 1.8, adjusted p-value: 0.02), and less likely to be synonymous nucleotide substitutions (OR: 0.6, adjusted p-value: 0.02). The identified eQTLs were more likely to be located in exons (OR: 1.8, adjusted p-value: 2.2×10^−12^), 5’UTRs (OR: 2.5 adjusted p-value: 4.2×10^−8^), 3’ UTRs (OR: 1.5, adjusted p-value: 1.6×10^−6^), and enhancers (OR: 1.8, adjusted p-value: 0.019), and less likely to be located in introns (OR: 0.9, adjusted p-value: 6.2×10^−10^) (Figure S4 D). Additionally, identified eQTLs are not significantly more likely to be either non-synonymous (OR: 1.1, adjusted p-value: 0.58) or synonymous nucleotide substitutions (OR: 0.9, adjusted p-value: 0.60).

The relationship between mRNA and protein abundance for genes with Bonferroni and FDR significant QTLs was, essentially, the same. Genes were defined in the same manner as the main text except using Bonferroni significance to define QTLs and are: 1) genes with both pQTLs and eQTLs (n=15); including 2) the subset of genes with sites that are both an eQTL and a pQTL (i.e. eQTL/pQTLs) (n=12); 3) genes with pQTLs but no eQTLs (n = 61); 4) genes with eQTLs but no pQTLs (n = 97); and, 5) all tested genes (n=5,739). Again, as we found in our main FDR analyses, genes with both eQTLs and pQTLs have a significantly higher correlation between mRNA and protein abundance than genes with eQTLs or pQTLs alone (Figure S4 E-F).

#### eQTL replication

We assessed the reproducibility of our eQTL results by comparing them with a previous eQTL meta-analysis of dorsolateral prefrontal cortex (dPFC) data from four cohorts by Sieberts *et al*. (*10*). We found a high degree of overlap between our eQTL results and the previously published results, which indicates that our methods for eQTL and pQTL identification are valid.

Sieberts *et al*. performed a meta-analysis using data from the dPFC of 1,433 cognitively normal and impaired individuals from the ROSMAP study, Mayo Brain Bank, Human Brain Collection Core, and the Common Mind Consortium. This study investigated proximal eQTLs within 1MB of an expressed gene, and declared eQTLs based on a False Discovery Rate (FDR) correction. Of the 35,064 eQTLs we identified, Sieberts *et al*. tested 31,340. Of these 31,340 sites, Sieberts *et al.* found 29,087 (92.8%) to also be eQTLs. Furthermore, for each SNV that either we or Sieberts *et al*. defined as an eQTL, we compared the sign and the magnitude of the test statistics. We found 81.9% (451,426 / 551,041) of the eQTL test statistics to have matching signs, and all eQTL test statistics to have a correlation of 0.77 between studies.

#### Relationship between Minor Allele Frequency and QTL identification

We examined the relationship between minor allele frequency (MAF) and our identified pQTLs and eQTLs by calculating the number and percentage of identified pQTLs (FDR < 0.05), eQTLs (FDR < 0.05), and eQTL/pQTLs that fall into the following MAF bins: 0.05 to 0.1, 0.1 to 0.2, 0.2 to 0.3, 0.3 to 0.4, and 0.4 to 0.5 (Table S5). We found that the percentage of the total pQTLs, eQTLs, and eQTL/pQTLs identified slightly increases with variant MAF. The smallest percentage of pQTLs, eQTLs and eQTL/pQTLs identified have a variant MAF between 0.05 and

0.1 (5%-8%), while the largest percentage of pQTLs, eQTLs, and eQTL/pQTLs have a variant MAF between 0.4 to 0.5 (26%-36%). We also examined the relationship between the MAF at each site in transcriptomic data (eQTL analysis) and the proteomic data (pQTL analysis), as these analyses had slightly different samples sizes (Figure S5). The correlation between the MAF of tall the sites in the pQTL and eQTL analyses is 0.98. Additionally, the correlation between the MAF in the pQTL and eQTL analyses at the sites identified as pQTLs, eQTLs, and pQTL/eQTLs, is 0.98, 0.98, and 0.97, respectively. These results suggest a slight, but not substantial, dependence of pQTLs and eQTLs on MAF.

#### Comparison of identified brain pQTLs with prior work in blood

We compared pQTLs in blood and brain by assessing the overlap between our pQTLs and pQTLs previously identified in blood by Emilsson et al. (*41*), Sun et al. (*42*) and Suhre et al. (*43*). The results of these three studies are available in the NHGRI-EBI Catalog of Published genome-wide association studies. Emilsson et al. used data from ~5,000 Icelanders over the age of 65, Sun et al. used data from 3,301 healthy blood donors from the INTERVAL study, and Suhre et al. used data from 1000 individuals from the KORA F4 study. All three studies used the SOMAscan platform to assay the plasma proteome.

The overlap between brain pQTLs and blood pQTLs was assessed by comparing our results to the results of Emilsson et al. (*41*), Sun et al. (*42*) and Suhre et al. (*43*). Together, these three studies report 3,014 unique blood pQTLs, of which 1,139 were available for testing in our brain (dorsolateral prefrontal cortex) data. This small overlap between the blood pQTLs and the sites that were tested in the brain indicates that the sets of proteins in brain and blood may be largely distinct. Of the 1,139 blood pQTLs also in our brain data, only 38 were found to be brain pQTLs (3%). These 38 blood and brain pQTLs are in 28 different genes.

---

**Fig. S1.**
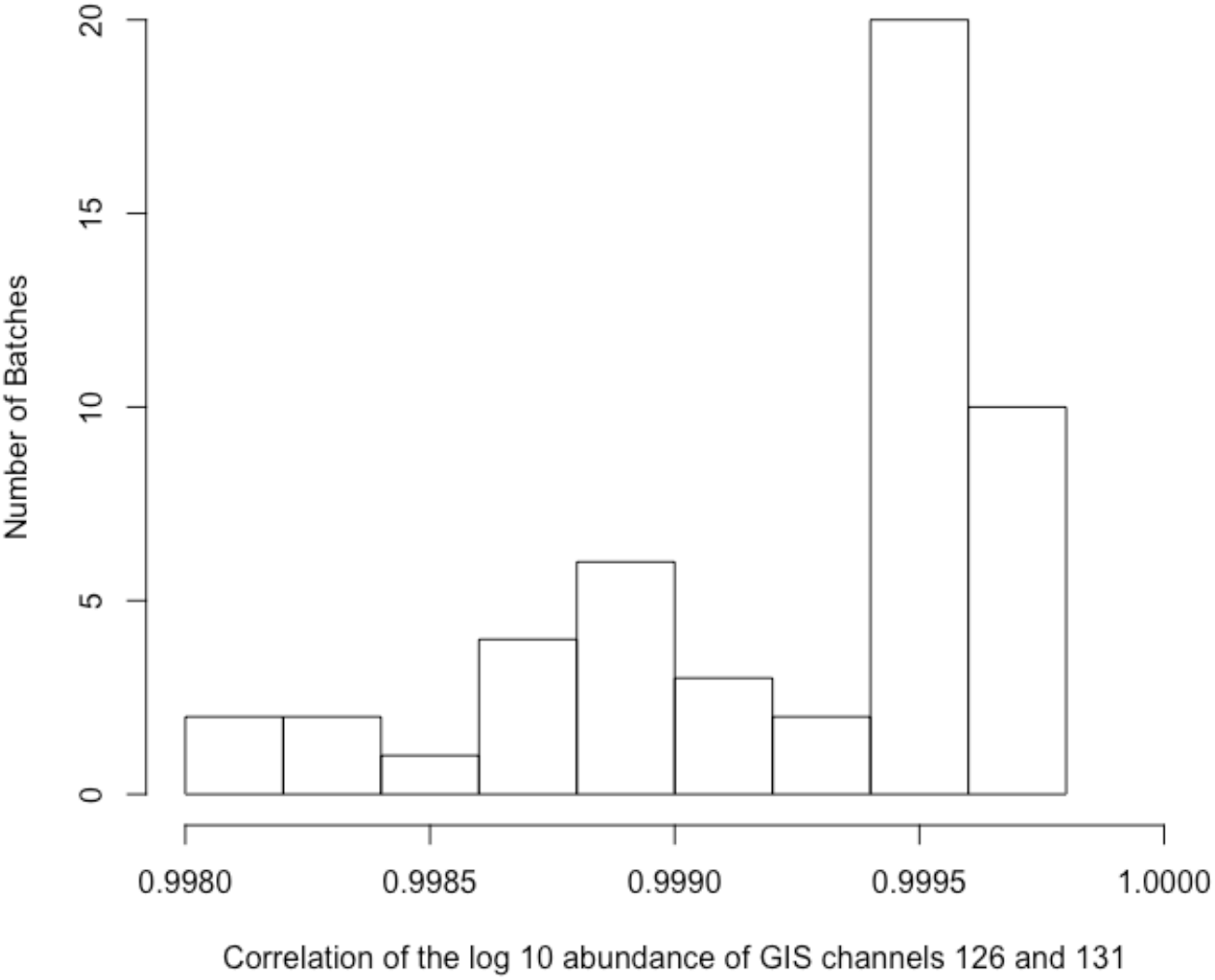
Distribution of the batch-specific correlation of GIS channels. Each TMT proteomic experiment, or batch, contains two GIS channels (126 and 131). Here we show the distribution of correlations of proteomic measurements between the batch-specific GIS channels.

**Fig. S2.**
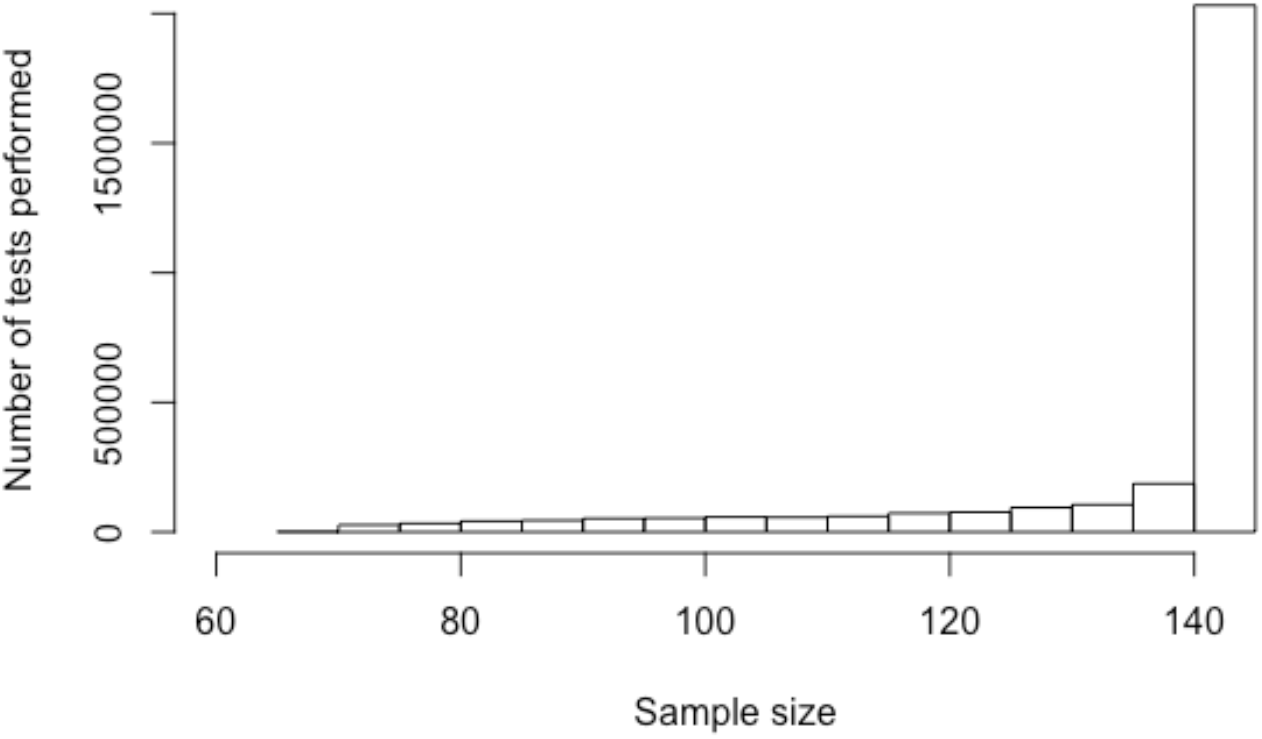
Distribution of the sample sizes used in the pQTL analyses. Due to the highly batch-specific nature of protein measurement in TMT proteomic experiments, each measured protein has a different sample size. This histogram shows the distribution of sample sizes across all regressions run to test the association between genotype and protein abundance for each SNV-protein pair.

**Fig. S3.**
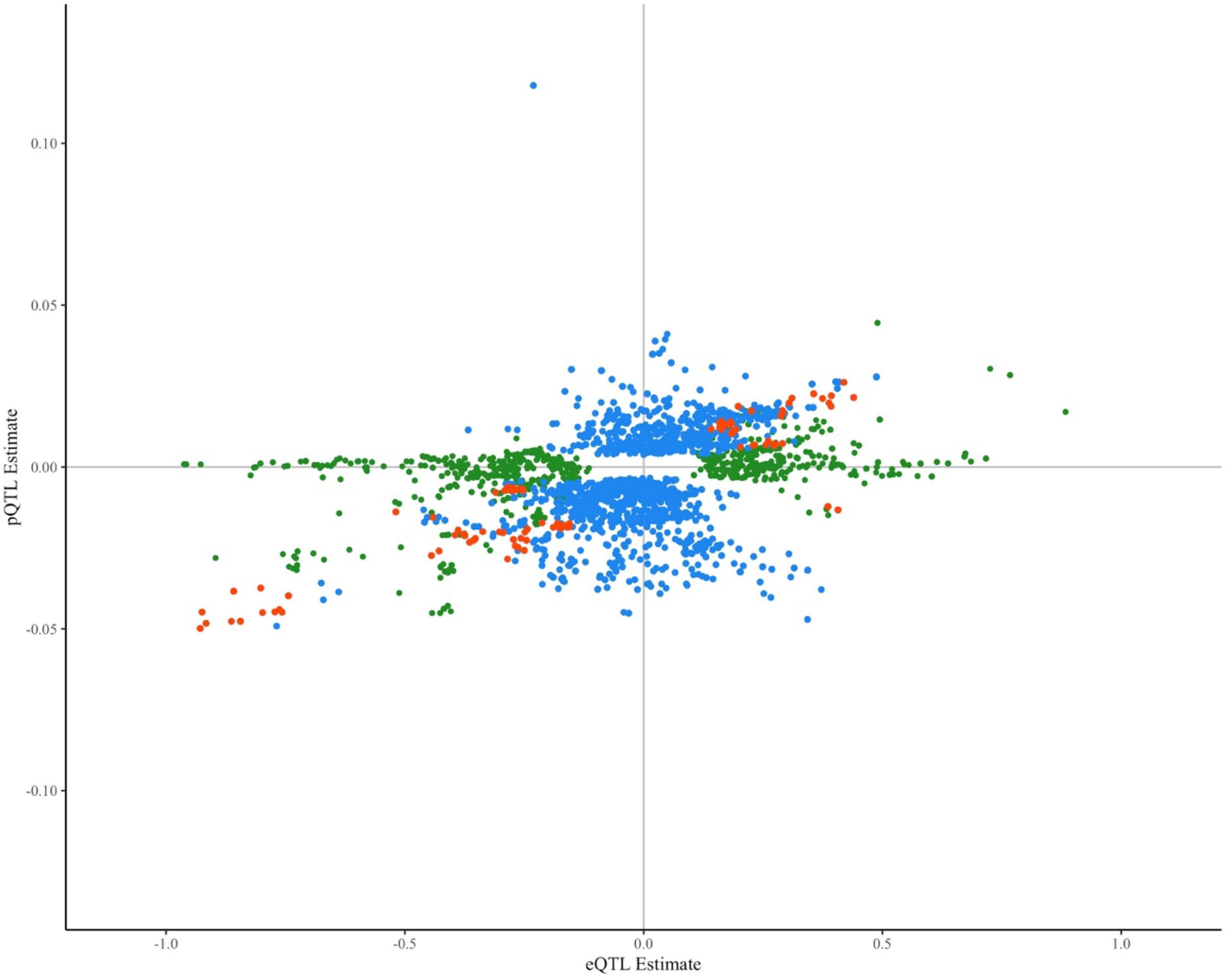
Comparison of eQTL and pQTL estimates for samples with paired transcriptomic and proteomic data. Each point represents one SNV tested against the abundance of the mRNA and protein of a single gene. eQTLs (defined based on FDR < 0.05) are shown in green, pQTLs (defined based on FDR < 0.05) are shown in blue, and sites that are both an eQTL and a pQTL (i.e. eQTL/pQTLs) are shown in orange.

**Fig. S4.**
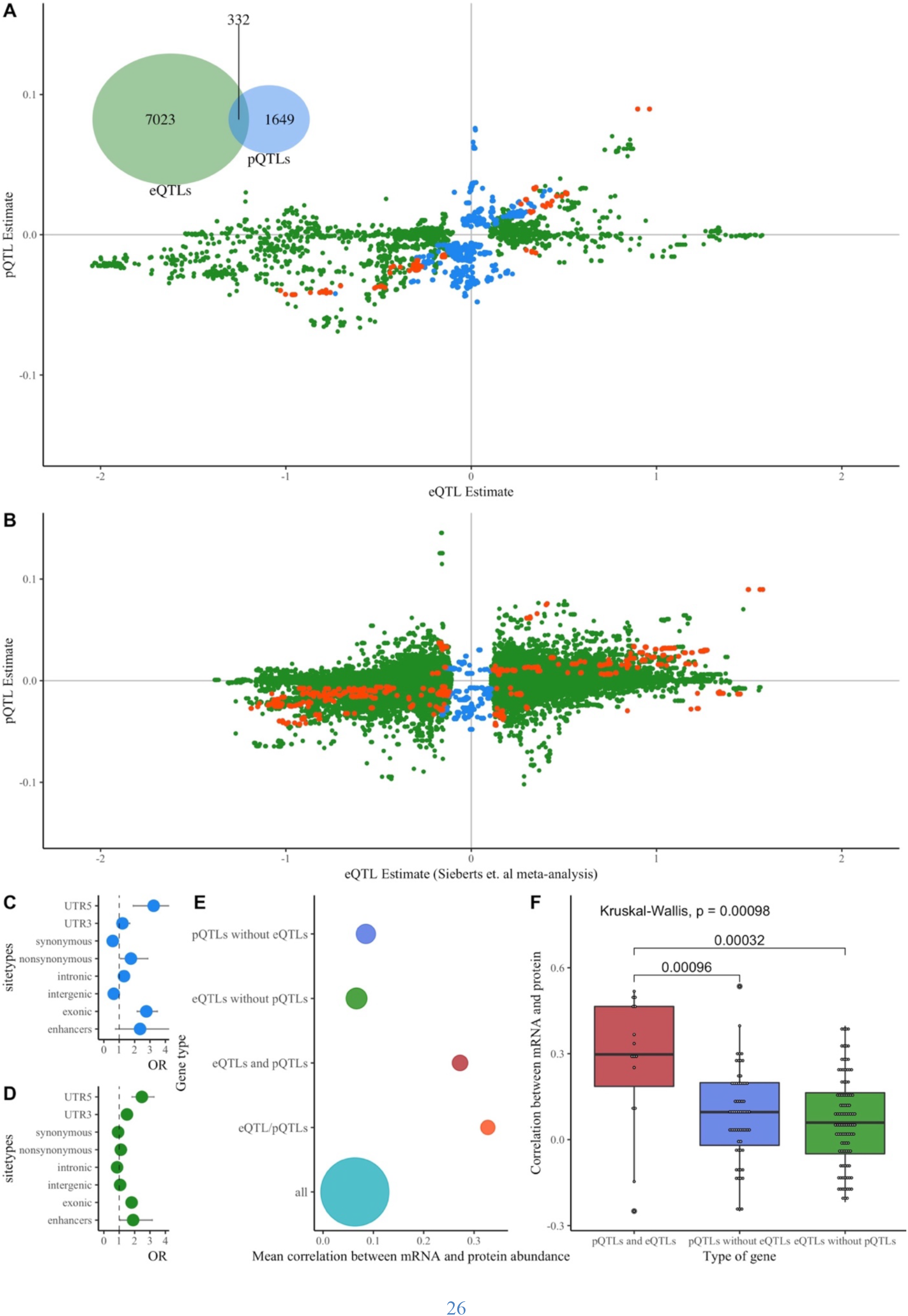
Protein and RNA Quantitative Locus Results Using a Bonferroni Significance Threshold. This figure summarizes the direction of effect and genomic annotation for pQTL and eQTL sites using QTLs defined based on Bonferroni significance threshold. (A) Comparison of eQTL and pQTL estimates. Each point represents one SNV tested against the abundance of the mRNA and protein of a single gene. eQTLs (defined based on Bonferroni correction < 0.05) are shown in green, pQTLs (defined based on Bonferroni correction < 0.05) are shown in blue, and sites that are both an eQTL and a pQTL (i.e. eQTL/pQTLs) are shown in orange. (B) Comparison of Sieberts *et al.* meta-analysis eQTL estimates (N=1433) and our pQTL estimates. Each point represents one SNV tested against the abundance of the mRNA and protein of a single gene. eQTLs (defined based on False Discovery Rate correction < 0.05) are shown in green, pQTLs (defined based on Bonferroni correction < 0.05) are shown in blue, and sites that are both an eQTL and a pQTL (i.e. eQTL/pQTLs) are shown in orange. (C) Results of Fischer’s exact tests assessing the overlap of pQTLs and genic locations. OR estimates are shown with 95% confidence intervals. (D) Results of Fischer’s exact tests assessing the overlap of eQTLs and genic locations. OR estimates are shown with 95% confidence intervals. (E) Mean correlation between mRNA and protein abundance for all genes, genes with eQTL/pQTLs (i.e. sites that are both an eQTL and a pQTL), genes with pQTLs and eQTLs, genes with pQTLs and no eQTLs, and genes with eQTLs and no pQTLs. The size of the point reflects the relative number of proteins within each gene type. (F) Comparison of genes with pQTLs and eQTLs, genes with pQTLs and no eQTLs, and genes with eQTLs and no pQTLs. P-values from the significant pairwise comparisons are shown.

**Fig. S5.**
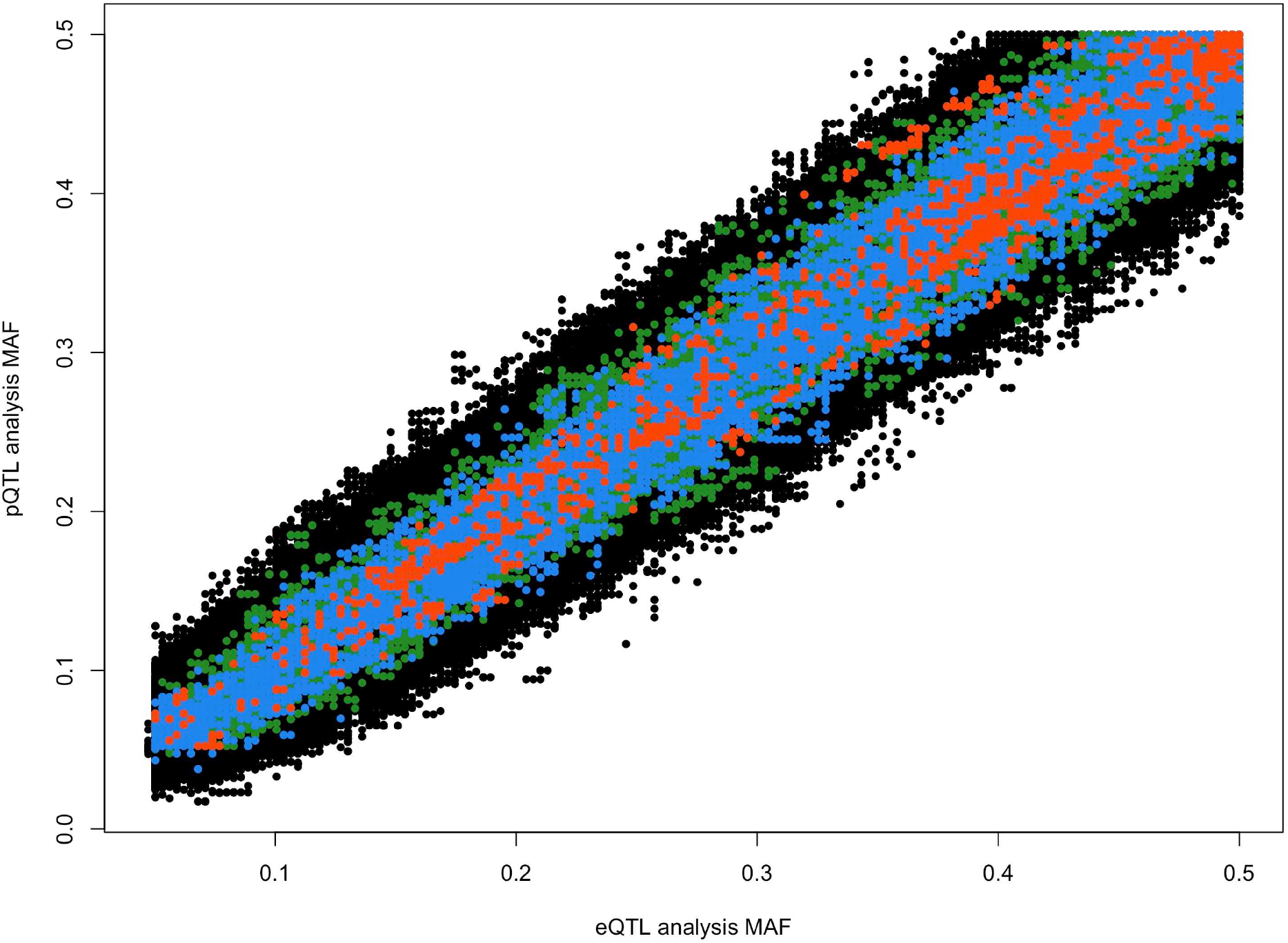
Comparison of the variant minor allele frequency for the eQTL and pQTL analyses. Each point represents one SNV tested against the abundance of the mRNA and protein of a single gene. eQTLs (defined based on FDR < 0.05) are shown in green, pQTLs (defined based on FDR < 0.05) are shown in blue, and sites that are both an eQTL and a pQTL (i.e. eQTL/pQTLs) are shown in orange. SNVs that are not a pQTL or eQTL are in black.

**Table S1.** Inflation in QTL analyses. We performed both pQTL and eQTL analyses on chromosome 1 with different combinations of covariates to assess their ability to reduce ***λ***, which estimates the amount of test statistic inflation. The covariates we assessed include: the first ten genetic principal components, the estimated cellular composition of each sample (i.e. the proportion of neurons, astrocytes, microglia, and oligodendrocytes), and the first five unknown confounders estimated using factor analysis.

| **pQTL analysis: chromosome 1** |  |
| --- | --- |
| Covariates | $\lambda$ |
| Genetic principal components (1 to 10) | 1.17 |
| Cellular composition (proportion of neurons, astrocytes, microglia, and oligodendrocytes) | 1.22 |
| Genetic principal components (1 to 10), cellular composition (proportion of neurons, astrocytes, microglia, and oligodendrocytes) | 1.17 |
| Genetic principal components (1 to 10), unknown confounders (5) | 1.17 |
| Genetic principal components (1 to 10), unknown confounders (5), cellular composition (proportion of neurons, astrocytes, microglia, and oligodendrocytes) | 1.18 |
| **eQTL analysis: chromosome 1** |  |
| Covariates | $\lambda$ |
| Genetic principal components (1 to 10) | 1.33 |
| Cellular composition (proportion of neurons, astrocytes, microglia, and oligodendrocytes) | 1.50 |
| Genetic principal components (1 to 10), cellular composition (proportion of neurons, astrocytes, microglia, and oligodendrocytes) | 1.49 |
| Genetic principal components (1 to 10), unknown confounders (5) | 1.70 |
| Genetic principal components (1 to 10), unknown confounders (5), cellular composition (proportion of neurons, astrocytes, microglia, and oligodendrocytes) | 1.78 |

**Table S2.**
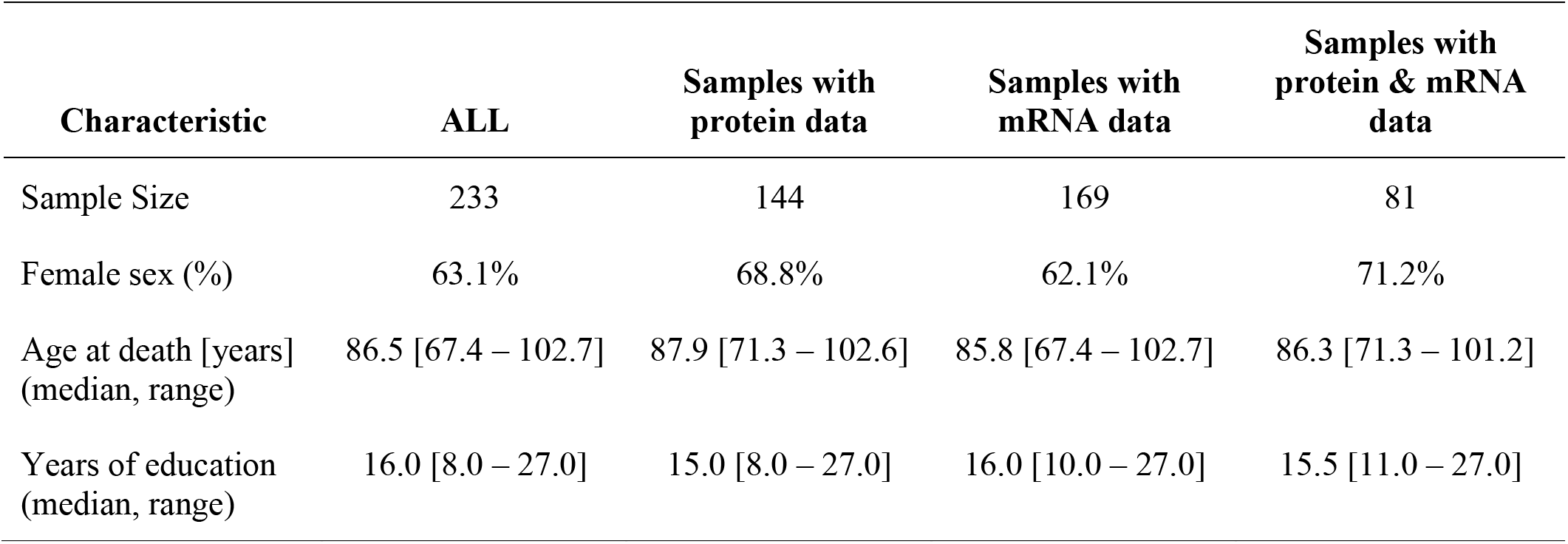
Demographics of analyzed ROS/MAP samples. All analyzed samples had a clinical diagnosis of no cognitive impairment at the time of death.

**Table S3.** Genes with sites that are both a pQTL and an eQTL (i.e. eQTL/pQTL). Both pQTLs and eQTLs were defined using a Bonferroni significance threshold of 0.05.

| Gene | Chr | # pQTLs | # eQTLs | #eQTL/pQTLs |
| --- | --- | --- | --- | --- |
| KYAT3 | 1 | 108 | 72 | 72 |
| GSTM5 | 1 | 23 | 52 | 22 |
| FAS | 10 | 36 | 25 | 25 |
| SNX32 | 11 | 27 | 4 | 4 |
| THEM4 | 1 | 80 | 140 | 80 |
| GSTM3 | 1 | 10 | 32 | 10 |
| GALM | 2 | 63 | 27 | 27 |
| PPIL3 | 2 | 48 | 56 | 48 |
| GSTT2B | 22 | 5 | 149 | 5 |
| TRPV2 | 17 | 2 | 30 | 2 |
| CAMLG | 5 | 19 | 19 | 19 |
| GPNMB | 7 | 18 | 18 | 18 |

**Table S4.** Top ten genes with the highest number of sites that are both an eQTL and a pQTL (i.e. eQTL/pQTLs). All the eQTLs and pQTLs were defined using a False Discovery Rate threshold of 0.05.

| Gene | Chr | # pQTLs | # eQTLs | #eQTL/pQTLs |
| --- | --- | --- | --- | --- |
| ERAP2 | 5 | 242 | 320 | 242 |
| GUF1 | 4 | 350 | 305 | 223 |
| NDUFAF1 | 15 | 159 | 172 | 153 |
| THEM4 | 1 | 139 | 161 | 139 |
| GPNMB | 7 | 136 | 139 | 136 |
| CSDC2 | 22 | 135 | 141 | 135 |
| GSTT2B | 22 | 134 | 212 | 133 |
| KYAT3 | 1 | 108 | 110 | 108 |
| GSTP1 | 11 | 141 | 113 | 105 |
| LARS2 | 3 | 108 | 166 | 105 |

**Table S5.** Relationship between minor allele frequency (MAF) and the number of pQTLs and eQTLs identified. All pQTLs and eQTLs were defined based on a False Discovery Rate threshold of 0.05.

| **MAF bin** | **# pQTLs (%) \| subset also eQTLs** | **# eQTLs (%) \| subset also pQTLs** |
| --- | --- | --- |
| *0.05 to 0.5* | *21,034 \| 3,364* | *35,064 \| 3,364* |
| 0.05 to 0.1 | 1,685 (8%) \| 163 (5%) | 2,640 (8%) \| 136 (5%) |
| 0.1 to 0.2 | 4,270 (20%) \| 568 (17%) | 6,845 (20%) \| 605 (18%) |
| 0.2 to 0.3 | 4,578 (22%) \| 549 (15%) | 7,528 (21%) \| 521 (15%) |
| 0.3 to 0.4 | 4,903 (23%) \| 778 (23%) | 8,542 (24%) \| 916 (27%) |
| 0.4 to 0.5 | 5,458 (26%) \| 1,213 (36%) | 9,381 (27%) \| 1,167 (35%) |

